# dbverse scales spatial omics analysis with embedded analytical databases

**DOI:** 10.64898/2026.08.09.743742

**Authors:** Edward C. Ruiz, Veronica Jarzabek, Jiaji Chen, Timur Rizvanov, Iqra Amin, Ruben Dries

## Abstract

Spatial omics datasets are increasing in size and complexity, exceeding the memory of standard computers and thereby limiting data analysis. Here we present dbverse, a framework for larger-than-memory matrix, spatial and genomic data analysis in embedded analytical databases. Benchmarks show dbverse provides orders of magnitude runtime improvements relative to established in-memory and file-backed methods for core operations in single-cell and spatial omics analysis. We integrated dbverse with Giotto Suite, scaling end-to-end preprocessing of millions of cells and enabling spatial alternative polyadenylation analysis as demonstrated on a Visium HD 3′ ovarian clear cell carcinoma sample. The dbverse framework provides an interoperable database foundation for larger-than-memory spatial omics analysis on ordinary computers.

## Main

Spatial omics technologies now profile molecular features *in situ* at increasing spatial resolution, molecular breadth, and cellular throughput^1–3^. Recent platforms can generate datasets with millions of cells per experiment, including cell-by-gene count matrices, cell and tissue geometries, and genomic interval data, each of which can reach hundreds of gigabytes per sample^4–12^. In parallel, experimental designs are shifting towards profiling multiple samples^13^ or exploring tissue architecture in 3D^14^ to achieve more robust statistical power^15^ or in-depth insights, respectively.

Together, this rapid growth in data size, formats, and scope of spatial omics projects has led to a growing memory footprint that now frequently exceeds the limits of commonly available computers (**Fig. 1a, Extended Data Table 1,** and **Methods**). Overcoming these limits typically requires migrating to expensive high-performance computing clusters with complex distributed frameworks, or relying on file-backed methods that can suffer from severe performance tradeoffs or limitations for more complex spatial data analysis workflows^16–18^. To circumvent these bottlenecks, researchers often resort to data simplification techniques, such as rasterization^19^, interpolation, or binning^20^, which improves computational efficiency but sacrifices native spatial resolution and limits downstream biological discovery. An ideal and practical solution should support larger-than-memory analysis at native resolution on ordinary computers while remaining compatible with established workflows.

**Figure 1.**
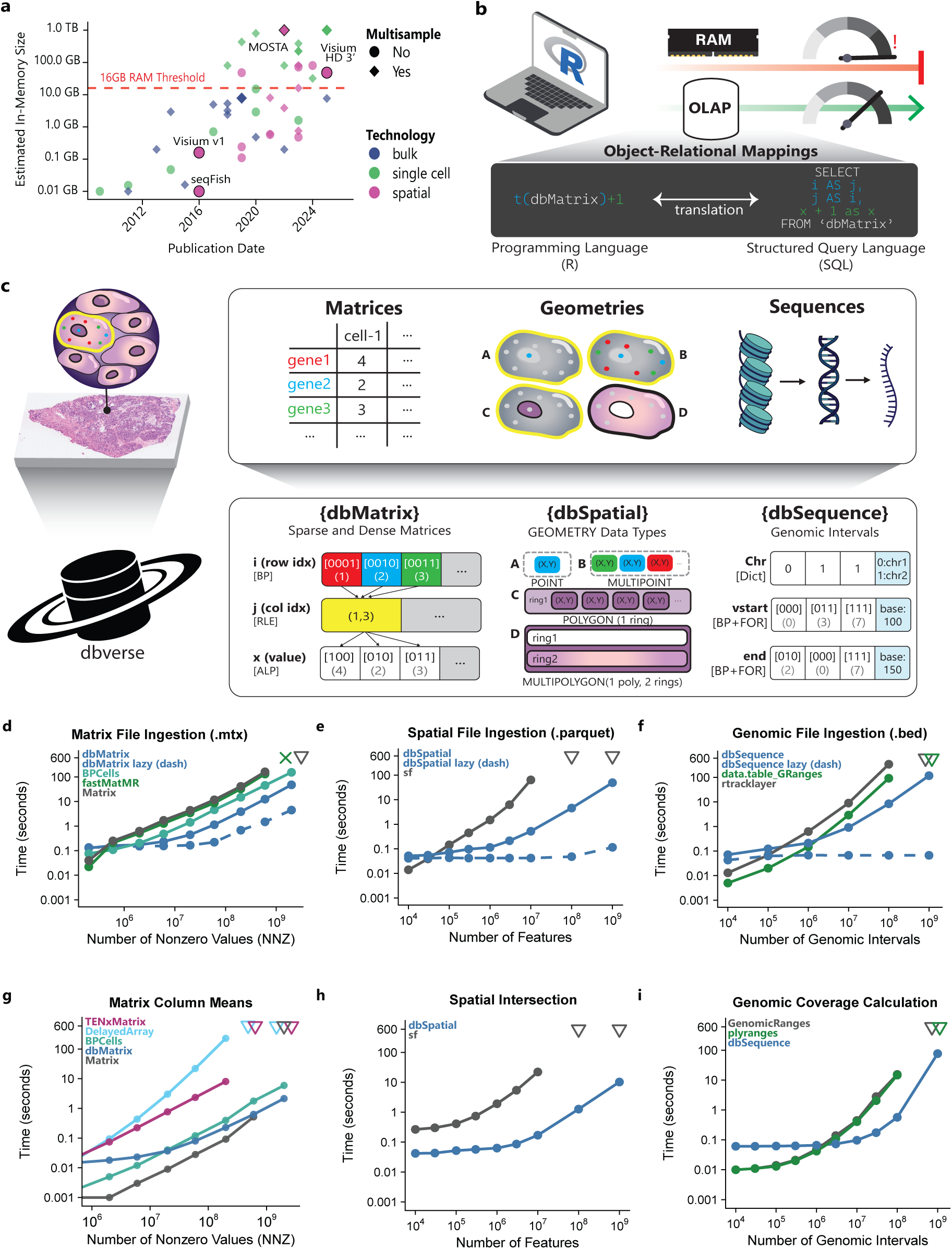
Overview of dbverse and quantitative benchmarks. **a,** Scatterplot depicting the size of various omics datasets relative to publication date. Circles represent single-sample datasets, diamonds represent multi-sample datasets; color indicates different omics technologies. The red dashed line marks a 16GB RAM threshold indicative of standard computers. **b,** Diagram showing the object–relational mapping paradigm of core analytical operations in R in the dbverse framework. **c,** Schematic of biological data types and their logical and physical representations in dbverse. dbMatrix, dbSpatial and dbSequence store matrices, geometries and genomic intervals, respectively, as compressed database tables with associated compression techniques. **d-f,** Line plots showing file ingestion runtime for Matrix Market (.mtx), GeoParquet point and BED genomic files. Colors indicate different packages while dashed lines denote lazy file registration. **g–i,** Line plots showing runtime benchmarks across increasing input sizes for core matrix, spatial, and genomic interval operations. Colors indicate different packages. ‘▽’ denotes a 600-s timeout or an operation not attempted after a preceding setup timeout; ‘×’ denotes an out-of-memory failure. Successful points show the median of five timed runs per method and dataset size.

To address these challenges, we developed dbverse, an analysis framework that executes common operations in single-cell and spatial omics data analysis directly within embedded or in-process online analytical processing (OLAP) databases^21,22^. The dbverse framework uses object–relational mappings (ORMs) to translate generic operations from common scientific packages into queries in Structured Query Language (SQL), which are executed by an embedded OLAP database (**Fig. 1b**). Currently, dbverse supports DuckDB^21^ and Apache DataFusion^22^ for specific operations in the framework (**Methods**). Specifically, we developed three standalone libraries for core omics data structures that are represented natively within the OLAP database: dbMatrix (sparse and dense matrices), dbSpatial (geometric features), and dbSequence (genomic intervals) (**Fig. 1c**). A fourth library, dbProject, manages database connections, defines base classes, and organizes shared helper functions across dbverse libraries (**Supplementary Fig. 1a**).

By delegating computations to OLAP databases, dbverse capitalizes on lazy evaluation and query optimization and larger-than-memory execution through disk spilling^23^. In addition, columnar compression techniques such as Run-Length Encoding (RLE)^24^, Bit Packing (BP)^25^, frame-of-reference (FOR), and Adaptive Lossless Floating-Point (ALP)^26^ further reduce storage footprint and read/write overhead. Interoperability with established scientific workflows is a core design principle of dbverse, which supports common scientific file formats, in-memory R objects, and the columnar binary Apache Arrow format^27^ (**Supplementary Table 1**). In addition, dbverse provides a multimodal API spanning over 80 matrix, 70 spatial, and 20 genomic operations (**Supplementary Table 2**). For supported methods, dbverse objects can replace conventional R objects with minimal changes to existing scripts due to their ORM design.

We evaluated dbverse against frequently used R packages in single-cell and spatial omics analysis. Specifically, we assessed both data ingestion times and key downstream operations on a commonly used computer with 16 gigabytes of system memory (**Methods**). Unlike the other evaluated methods, dbverse supports both eager and lazy data registration for all tested file types. The lazy path registers files without immediately materializing them in the database, allowing subsequent multi-step workflows to be optimized as a single query plan. For Matrix Market (.mtx) files, dbMatrix was the only method evaluated offering lazy registration. At two billion nonzero matrix entries, eager dbMatrix was 3× faster than BPCells^28^, whereas Matrix^29^ exceeded 600 seconds and fastMatMR^30^ encountered an out-of-memory error (**Fig. 1d**). For spatial geometry data, dbSpatial was the only evaluated method to complete eager ingestion of one billion GeoParquet point features, whereas sf^31,32^ failed to complete within the time window at 100 million and one billion point features (**Fig. 1e**). Finally, lazy dbSequence ingestion registered up to one billion genomic BED intervals, and the eager dbSequence path was the only method to complete within the timeout window at that scale. At 100 million intervals, eager dbSequence was approximately 34× faster than rtracklayer^33^ and approximately 10× faster than the combined data.table and GenomicRanges^34^ path (**Fig. 1f**). Next, we benchmarked common matrix, spatial and genomic operations and showed representative examples. Compared to other in-memory methods such as Matrix^29^ and DelayedArray^35^, and on-disk methods such as the sparse HDF5-backed TENxMatrix^36^ and BPCells^30^, dbMatrix was the fastest to calculate column means at the largest tested scale, approximately 3× faster than BPCells^28^, and completed the operation within our time limit at every input size (**Fig. 1g**). Supporting benchmarks further showed that dbMatrix maintained compact on-disk storage. At the largest scale, BPCells^28^ used 3.4× more storage and dense HDF5Matrix^36^ used 1.9× more storage than dbMatrix (**Supplementary Fig. 1b**). dbMatrix was also fastest at the largest successfully evaluated random row-and-column indexing scale (**Supplementary Fig. 1c**). For spatial operations, dbSpatial was the only evaluated method that completed the counting of points intersecting at least one polygon for one billion input points within the time limit (**Fig. 1h**). For genomic operations, only dbSequence could be evaluated at one billion intervals within the time limit. At 100 million intervals, dbSequence was approximately 31× faster than plyranges^37^ and 34× faster than GenomicRanges^34^ (**Fig. 1i**). Extended benchmarks showed similar performance advantages across additional matrix, spatial and genomic operations among method–operation combinations that completed successfully (**Supplementary Table 3**).

To determine whether the dbverse R packages could scale an established spatial-omics analysis ecosystem, we integrated them into Giotto Suite^38^. We developed GiottoDB, an R package that extends the Giotto object by replacing in-memory expression matrices and spatial geometries with dbMatrix and dbSpatial objects, respectively, thereby enabling database optimizations like query pushdown for spatial omics data analysis (**Fig. 2a**). We also developed GiottoSeq, which adds alternative polyadenylation (APA) analysis to Giotto by supporting outputs from scAPAtrap^39^ and movAPA^40^ and using dbSequence and dbMatrix operations for scalable poly(A)-site aggregation and annotation (**Fig. 2b** and **Supplementary Fig. 1d**). Together, these packages provide a database-backed workflow for joint spatial analysis of gene expression and poly(A)-site usage.

**Figure 2.**
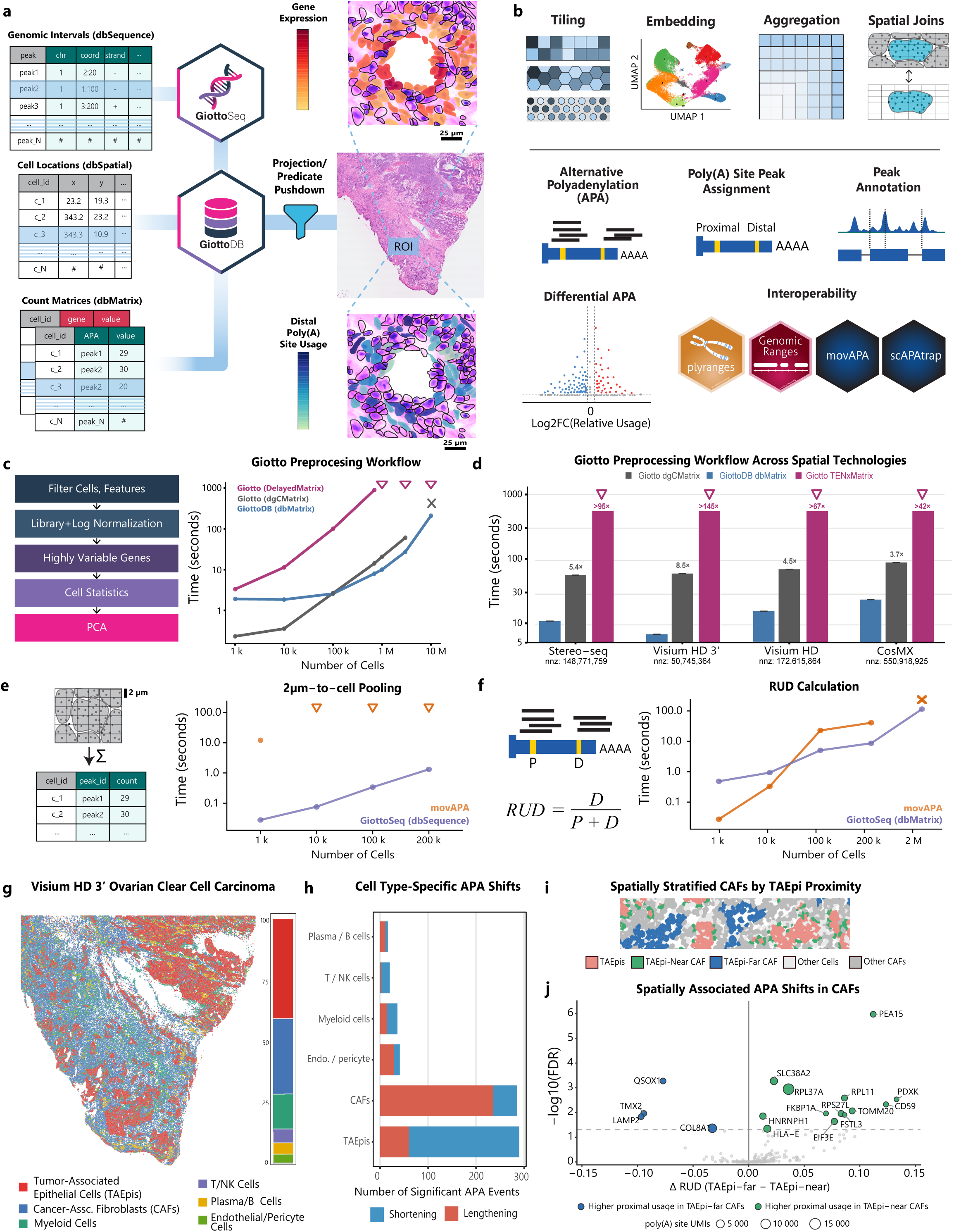
dbverse and Giotto Suite integration to scale spatial workflows and enable spatial APA analysis. **a,** Diagram showing the organization and workflows for dbverse-backed GiottoDB and GiottoSeq packages. Images show gene expression and distal poly(A)-site usage. Scale bars, 25 μm. **b,** GiottoSeq functions for spatial processing, APA analysis and interoperability with R and Bioconductor packages. **c,** Diagram and line plot showing end-to-end Giotto preprocessing workflow benchmark across increasing semisynthetic dataset sizes. ‘▽’ denotes a 1,000-s timeout or a larger input not run after a preceding timeout and ‘×’ denotes out-of-memory failure. Successful points show the median of five runs per method and dataset size. **d,** Barplot with error bars showing median preprocessing workflow runtimes across spatial technologies with Giotto and GiottoDB. ‘▽’ denotes runs exceeding 1,000 seconds and ‘×’ denotes out-of-memory failure. Successful bars show the median of five runs per method and dataset, errors are shown. ‘nnz’ indicates the number of non-zero matrix elements. **e,** Diagram and line plot showing benchmark runtimes for 2-µm bin-to-cell pooling. ‘▽’ denotes a 600-s timeout or a larger input not run after a preceding timeout. Successful points show the median of five runs per method and dataset size. **f,** Line plot showing runtimes for calculating relative usage of distal poly(A) sites (RUD). ‘×’ denotes an out-of-memory failure. Successful points show the median of five runs per method and dataset size. **g,** Spatial map and corresponding barplot depicting cell types and tissue composition, respectively. **h,** Horizontal barplots showing differential APA events by cell type (one-vs-rest). Colors represent 3’UTR shortening or lengthening as indicated. **i,** Spatial region of interest showing cancer-associated fibroblasts (CAFs) stratified by distance to neighboring tumor-associated epithelial cells (TAEpis). TAEpi-near CAFs, TAEpi-far CAFs, other CAFs, TAEpis and other cells shown separately. **j,** Volcano plot depicting TAEpi-proximity-associated APA shifts in CAFs. Colored points have FDR < 0.05; color indicates the spatial context with higher proximal poly(A)-site usage.

We applied this framework to a Visium HD 3′ ovarian clear cell carcinoma (OCCC) dataset containing gene-expression and 3’-end sequence information at 2µm resolution. We first assessed whether GiottoDB could scale standard preprocessing operations required for downstream analysis, including backend setup, filtering, normalization, feature selection and dimensionality reduction. Across semisynthetic datasets (**Methods**), GiottoDB completed the end-to-end workflow through 10 million cells. Giotto with the default in-memory dgCMatrix backend completed the workflow through 3 million cells but encountered an out-of-memory failure at 10 million cells. The sparse HDF5-backed TENxMatrix path was approximately 39× slower than GiottoDB at 100,000 cells and exceeded the 1,000-second time limit at one million cells and larger (**Fig. 2c**). Across four spatial-omics technologies, GiottoDB was up to 8.5× faster than the dgCMatrix workflow, whereas the TENxMatrix workflow exceeded 1,000 seconds for every real-world dataset^4,6,9^ (**Fig. 2d**).

Having established end-to-end workflow scalability with GiottoDB, we next evaluated two APA-specific bottlenecks in the OCCC dataset: pooling peak-by-2-µm-bin counts into a peak-by-cell matrix and calculating the relative usage of distal poly(A) sites (RUD). GiottoSeq with dbSequence completed bin-to-cell pooling at every tested scale, whereas the movAPA implementation completed only the 1,000-cell dataset and exceeded 600 seconds at larger sizes (**Fig. 2e**). For RUD calculation, movAPA was faster at 1,000 and 10,000 cells, whereas GiottoSeq was approximately 2.5× faster at 100,000 cells and 3.1× faster for the full 200,769-cell dataset. At 2.1 million cells, GiottoSeq completed the calculation, whereas movAPA encountered an out-of-memory failure (**Fig. 2f**).

These scalable operations enabled us to examine whether the dataset could resolve cell-type-specific and spatially associated differences in poly(A)-site usage. We first identified six broad cell-type groups in the Visium HD 3′ OCCC dataset (**Fig. 2g** and **Supplementary Fig. 1e**). Tumor-associated epithelial cells (TAEpis) showed the largest number of candidate differential APA events, with a bias toward increased proximal poly(A)-site usage, consistent with the widespread 3’UTR shortening previously reported in malignant cells^41^. Cancer-associated fibroblasts (CAFs), in contrast, showed a relative bias towards increased distal poly(A)-site usage in this sample (**Fig. 2h**). Consistent with the coverage dependence reported in previous single-cell APA analyses^42^, cell-type comparisons with greater combined poly(A)-site UMI depth tended to yield more candidate differential APA events in our dataset (**Supplementary Fig. 1f-h**). Finally, to evaluate whether we could identify spatially associated APA shifts, we stratified CAFs by proximity to neighboring TAEpis (**Methods** and **Fig. 2i**) and identified candidate APA shifts between TAEpi-near and TAEpi-far CAFs, including genes with higher proximal-site usage in either spatial context (**Fig. 2j**). These findings suggest that local TAEpi–CAF context may be associated with variation in poly(A)-site usage.

In conclusion, dbverse provides a database-backed foundation for larger-than-memory single-cell and spatial omics analysis in R. By translating operations from established matrix, spatial and genomic-interval APIs into queries executed by embedded OLAP databases^21,22^, dbverse enables scalable analyses on ordinary computers while supporting interoperability with common R objects, file formats and supported analysis workflows. Through GiottoDB and GiottoSeq, we show that this architecture can extend established spatial omics frameworks to full-resolution, multimodal datasets and support analyses that would otherwise be limited by memory or file-backed performance bottlenecks. More broadly, these results demonstrate the performance benefits of embedded analytical databases for emerging high-resolution omics workflows that exceed system memory on widely available computers.

## Methods

### Description of dbMatrix library

The dbMatrix library supports over eighty composable matrix operations (**Supplementary Table 2**) within an embedded OLAP database (DuckDB)^21^. Sparse and dense matrices are represented as a database table in the coordinate list (COO) format (**Fig. 1c**). dbMatrix represents sparse (dbSparseMatrix) and dense (dbDenseMatrix) matrices, with the latter supporting inclusion of zero-valued entries (dense COO tables). The schema of the sparse and dense COO table consists of three columns: *i* (row index, data type: int32 or int64), *j* (column index, data type: int32 or int64), and *x* (element value, data type: double). DuckDB^21^ applies lightweight per-column compression on this schema, using algorithms such as Bit-Packing (BP) for row indices^25^, Run-Length Encoding (RLE) for column indices^24^, and Adaptive Lossless Floating-Point (ALP) compression^26^ or RLE for matrix values (**Fig. 1c**).

Implemented in the S4 class system in R, the dbMatrix R package emulates select classes and methods from the Matrix R package^29^, containing attributes (slots) held in memory for name (character), value (pointer to COO table in the database^21^), dims (an integer vector of length two), and dim_names (list of length two containing row and column names) (**Supplementary Fig. 1a**). It supports nearly all S4 group generic functions for performing arithmetic (“+”, “−”, “*”, “^”, “%%”, “%/%”, “/”), comparisons (“==”, “>”, “<”, “!=”, “<=”, “>=”), logical expressions (“&”, “|”), mathematical functions (“abs”, “sign”, “sqrt”, “ceiling”, among others), and summary operations (“max”, “min”, “prod”, “sum”, “any”, “all”) between dbMatrix objects, numerical vectors, scalar values, and special numeric values (e.g. −Inf, Inf, NaN, 0, NA).

### Description of dbSpatial library

The dbSpatial library provides access to over 160 spatial operations (**Supplementary Table 2**) as R bindings to C++ functions in the DuckDB spatial extension. The dbSpatial R API supports over twenty generics from the sf R package^31,32^. S4 dbSpatial objects represent database tables where at least one column utilizes a GEOMETRY data type, adhering to the simple features specification^43^ to encode established geometry subtypes (**Fig. 1c**). Objects from the dbSpatial R package can also be efficiently converted to terra^44^ or sf ^31,32^ objects with support for reading and writing to/from more than fifty spatial file formats (**Supplementary Table 1**).

### Description of dbSequence library

The dbSequence library is designed for the querying and manipulation of genomic intervals. Genomic interval data is represented in the database utilizing a schema for chromosome, start position, and end position. For BED files, this structure is compressed via Dictionary ([Dict]) encoding for repetitive chromosome identifiers, and a combination of Bit-Packing and frame-of-reference ([BP+FOR]) encoding for integer genomic coordinates (**Fig. 1c**). This architecture allows for the efficient execution of genomic interval overlaps, peak-to-gene aggregation, and sequence coverage calculations at scales that exceed the memory limitations of alternative methods. Like other dbverse packages, dbSequence also supports over twenty database-backed methods from common Bioconductor packages (**Supplementary Table 2**) and conversion to and from GRanges^34^ objects. The dbSequence package is also compatible with the exonR package^45^, which supports ingestion and CRUD operations for genomic file formats in Apache DataFusion^22^.

### Database connection and state management via dbProject

The dbProject library serves as a lightweight utility package for managing base classes, utility functions, and database management across dbverse R packages (**Supplementary Table 2**). The dbProject package defines the dbData virtual S4 base class (containing slots ‘name’ and ‘value’), which underlies all R classes in dbverse (**Supplementary Fig. 1a**). To address the session-bound nature of database connections in R, dbProject implements an optional R6 class which serves as a mutable object for managing dbverse objects. This R6 object also centralizes database directory mapping, ensuring automatic, lock-safe reconnections when sessions drop or restart. To enable project and dbData object persistence, dbProject integrates with the pins R package^46^. This allows researchers to safely serialize, track, and restore large database objects, lazy SQL queries, and metadata representations via a persistent, version-controlled manifest. In effect, complex workflows consisting of in-memory and database objects can be paused, shared, and resumed without recomputing expensive intermediate steps or losing underlying data references across sessions.

### Scaling spatial omics analysis via GiottoDB

To demonstrate the real-world benefit of dbverse for end-to-end spatial omics analysis, we developed GiottoDB, a package that bridges methods from dbverse with the Giotto Suite^47^ ecosystem. The GiottoDB package defines a new GiottoDB S4 object which inherits from the giotto object and incorporates a connection slot while maintaining API compatibility with many functions from the Giotto Suite ecosystem (**Supplementary Fig. 1d**). GiottoDB replaces memory-bound slots in the giotto object with database representations from dbverse, substituting in-memory or file-backed matrices with dbMatrix objects in the expression slots, and replacing SpatVector geometries from the terra^44^ R package with dbSpatial objects in the spatial_info and feat_info slots. Notably, because dbverse objects emulate standard R classes and their APIs, within supported workflows, GiottoDB objects can be used with existing Giotto code with minimal code modifications. To achieve this, we implemented S3 and S4 methods within GiottoDB that reroute standard Giotto functions to database-backed dbverse implementations when necessary. For example, specific downstream operations that strictly require in-memory structures (e.g. differential expression testing or plotting via ggplot2) contain GiottoDB methods, which employ lazy-evaluated materialization. This strategy safely casts only the strictly required data subsets into RAM while leaving the parent object securely backed by the database with memory thresholds configurable by global options. Finally, custom saveGiotto and loadGiotto methods safely package the embedded database file alongside the standard giotto output directory, utilizing dbData metadata to automatically reconnect stale database pointers.

### Polyadenylation Site Identification and Filtering

To define poly(A) sites for downstream APA analysis in the OCCC Visium HD 3’ dataset, initial poly(A) site peaks and counts were identified using scAPAtrap^39^ from the raw spatial sequencing data (**Code Availability**). The resulting peak outputs were imported into the movAPA^40^ R package to construct a PACdataset object. False-positive peaks originating from internal priming artifacts were computationally removed using the UCSC hg38 human reference genome (movAPA::removePACdsIP, returnBoth = TRUE, up = −10, dn = 10, conA = 6, sepA = 7), and only the retained non-internal-priming sites were used for downstream analysis. The remaining poly(A) sites were mapped to genomic features using the TxDb.Hsapiens.UCSC.hg38.knownGene annotation database. Rather than extending 3’ untranslated regions (3’UTRs), sites were restricted to annotated 3’UTR intervals from GenomicFeatures::threeUTRsByTranscript using strand-aware overlaps. The resulting strict 3’UTR PACdataset, retained without additional count-based filtering prior to pooling, contained 22,259 poly(A) sites across 11,299 genes and 7,129,832 2µm spatial bins, with 31,696,449 poly(A) site UMIs, and was used as the reference poly(A) site dataset for downstream spatial aggregation.

### Visium HD 3’ Ovarian Cancer Dataset Processing and Clustering

A GiottoDB object was generated from the Visium HD 3′ dataset. Entries below an expression threshold of 1 were filtered, features detected in fewer than three cells were removed, and cells with fewer than 50 detected features were excluded. Cells with zero RNA library size after filtering were removed before normalization. Read counts were normalized using a library-size scaling factor of 10,000 and log2-transformed (offset = 1). Highly variable features were identified using a LOESS fit on the coefficient of variation. Dimensionality reduction was performed with the runPCA function to compute 20 centered and scaled principal components. UMAP embeddings were computed from principal components 2–20 (n_neighbors = 30, min_dist = 0.3, metric = cosine, n_epochs = 400, seed = 1234). A shared nearest-neighbor graph was then constructed from principal components 2–20 (k = 30, minimum shared = 5, top shared = 3) and Leiden clustering was performed (resolution = 0.2, modularity objective, 1000 iterations; seed = 1234). Clusters were assigned to six broad biological cell-type groups plus a low-quality/unclear category using canonical marker genes (**Supplementary Fig. 1e**). These broad annotations were then used for spatial visualization, cell-type aggregation and differential APA summaries.

### Spatial Alternative Polyadenylation and Proximity Analysis with GiottoSeq and GiottoDB

To profile alternative polyadenylation (APA) in Visium HD 3’ data, poly(A) site counts from 2µm bins were joined to segmented-cell barcode mappings to assign cell IDs, pooled by peak and cell using dbSequence::pool(), converted to a dbMatrix, and aligned to the RNA matrix on shared cell IDs. Peak metadata from the strict 3’UTR PACdataset were attached to the GiottoDB object as a GiottoSeq peaksObj using createpeaksObj() and setPeaks(). With GiottoSeq, proximal and distal poly(A) site pairs were identified, requiring a genomic distance between 50 and 10,000 base pairs and a minority usage ratio of ≥0.01 (minRatio = 0.01). Relative usage of distal sites was computed at both the single-cell and aggregated cell-type levels using the smartRUD method as previously described^40^. Cell-type aggregation was performed with GiottoSeq::aggregateCells(by = “giotto_cell_type”). Differential APA between cell types (one vs all) was assessed using Fisher’s exact tests on proximal and distal counts with min_counts = 20, and Benjamini-Hochberg correction. The low-quality/unclear cell type category was excluded from differential summaries. To summarize APA shifts in the sparse Visium HD 3′ data following cell-type aggregation, we treated events passing an exploratory threshold (adjusted P < 0.10, absolute log2 fold change > 0.10) as candidate APA events. Positive log2 fold change denotes 3’UTR lengthening and negative log2 fold change denotes 3’UTR shortening.

To evaluate spatially organized APA in the OCCC sample, CAFs and tumor-associated epithelial cells (TAEpis) were identified from the cell-type annotations. For each CAF, the Euclidean distance from its segmented-cell centroid to the nearest TAEpi centroid was calculated. Using the Space Ranger image scale of 0.273706 µm per full-resolution pixel, CAFs within 13.7 µm (≤50 pixels) of a TAEpi were classified as TAEpi-near, whereas CAFs at least 54.7 µm (≥200 pixels) away were classified as TAEpi-far. Intermediate CAFs were shown in the spatial map but excluded from the near-versus-far differential APA test. For each distance group, poly(A)-site counts were pseudobulked by gene and proximal/distal poly(A)-site assignment. TAEpi-proximity-associated APA shifts were tested per gene using Fisher’s exact tests on distal and proximal poly(A)-site counts, with Benjamini-Hochberg correction. RUD was calculated for each group, and ΔRUD was defined as RUD in TAEpi-far CAFs minus RUD in TAEpi-near CAFs. Candidate genes required at least 20 poly(A)-site UMIs in each group and at least 100 total poly(A)-site UMIs for testing. Figure 2j shows genes with at least 1,000 total poly(A)-site UMIs; colored and labeled points have FDR < 0.05. Positive ΔRUD indicates higher distal-site usage in TAEpi-far CAFs and therefore higher proximal-site usage in TAEpi-near CAFs, with the converse for negative ΔRUD. These tests describe within-section count differences, and analysis of a single tissue section does not assess between-sample reproducibility.

### Estimation of omics dataset sizes

Published omics dataset dimensions were compiled from the sources listed in **Extended Data Table 1**. To compare published omics dataset sizes in **Fig. 1a**, we estimated the size of a fully materialized dense matrix as the number of measured features multiplied by the number of observations and 8 bytes per entry (double-precision numeric values in R), divided by 10^9^ to obtain decimal gigabytes. Observations represent cells, spatial units or samples, as appropriate. Size estimates do not include other raw data modalities, metadata, or intermediate copies whose contributions vary by dataset.

### Synthetic and semisynthetic benchmark data generation

Figure 1d,g and Supplementary Figure 1b,c used deterministically generated 20,000-feature sparse matrices with increasing column counts and fixed 99% sparsity. Figure 1e,h used deterministically generated GeoParquet point and polygon geometry datasets. Figure 1f,i used deterministically generated GRCh38-scale BED intervals and each smaller BED file was a subset of the one-billion-interval file. For Figure 2c, semisynthetic count matrices containing 2,000 genes and between 1,000 and 10 million columns were generated from the Visium HD CytAssist colorectal cancer 8-µm-bin matrix. Genes were sampled across the observed range of detection frequencies in the original matrix, and additional matrices were generated by resampling while retaining library-depth structure. For Figure 2f, a 2.1-million-cell semisynthetic APA dataset was generated from the OCCC data by resampling observed profiles within cell-annotation and library-depth groups. These semisynthetic datasets were used only to evaluate computational scaling. Biological analyses in Figure 2g-j were based on the original OCCC dataset.

### Benchmarking dbverse libraries

Reproducible R scripts benchmarking each library are provided (**Code Availability**). Each successful method–size combination included one untimed warm-up followed by five timed replicates, with garbage collection before each timed replicate, on a Mac mini (16 GB RAM, 256 GB SSD, M4 processor) with a 1 TB Thunderbolt 4 NVMe external solid-state drive (Samsung 990 PRO). For Matrix Market (.mtx) ingestion, dbMatrix was benchmarked against Matrix^29^, fastMatMR^30^, BPCells^28^. For column-mean calculation, dbMatrix was compared with Matrix^29^, BPCells^28^, DelayedArray^35^ and the sparse HDF5-backed TENxMatrix^36^. dbSpatial was benchmarked against sf^31,32^ for GeoParquet point ingestion and point-in-polygon counting. We benchmarked dbSequence against rtracklayer^33^ and a combined data.table and GenomicRanges^34^ path for BED ingestion, and against GenomicRanges^34^ and plyranges^37^ for binned coverage computation. Figure 1g–i timed only the reported operation. Backend setup had its own 600 second timeout limit. Figure 2c,d benchmarks used a 1,000 second per-replicate timeout. Figures 1g-i, and 2c,d were run in isolated processes with a 14GiB RSS and 64GB R vector limit. When a method–size combination encountered its first timeout or out-of-memory failure, subsequent planned replicates were assigned the same classification without being re-executed. Median times therefore describe combinations with five successful timed replicates.

Figure 1d,e,g,h, Supplementary Figure 1b,c, the matrix and spatial Supplementary Table 3 benchmarks, and Figure 2c,d were run under R 4.5.1, Bioconductor 3.22.0 and DuckDB 1.5.2. Figure 1f and 1i, the benchmarks in Supplementary Table 3, and Figure 2e and 2f were run in a separate dbSequence environment (R 4.6.1, Bioconductor 3.23.1, DuckDB 1.5.4.3), with package versions recorded in the accompanying renv.lock files (**Code Availability**). For Figure 1h, we counted distinct points intersecting at least one of 100,000 fixed grid polygons, while Figure 1i measured coverage over fixed 1 kb bins spanning chr1:1–1,000,000. Where methods completed successfully, outputs were checked for equivalence. Supplementary Figure 1b reports on-disk size in bytes for the dbMatrix, BPCells, dense HDF5Matrix, and sparse TENxMatrix representations of identical inputs. All methods ran in isolated sequential workers, and successful results agreed within an absolute tolerance of 1e^−8^. Supplementary Table 3 also includes a polygon–polygon intersection benchmark between equal-sized grid and irregular polygon layers. Setup costs were measured separately and excluded from that table. Where benchmarks overlapped with Figure 1g–i, we reused the finalized measurements and validated successful results against the reference methods.

The workflow benchmark in Figure 2c and 2d timed backend setup, filtering, normalization, highly variable feature selection, principal component analysis, and cell-level summary-statistic calculation. Data download, input preparation, and format conversion all took place before timing began. Filtering applied expression_threshold = 0, feat_det_in_min_cells = 1, and min_det_feats_per_cell = 1. Normalization used a 10,000 scale factor with log transformation and no feature or cell scaling. For highly variable features, we used method = “var_p_resid” with var_number = 1,000, and PCA used 20 components with centering but no explicit unit-variance scaling; addStatistics computed cell statistics only. Each backend was given a requested budget of eight cores. Because Giotto’s public dgCMatrix and TENxMatrix paths effectively run under SerialParam(1) via BiocParallel, they ran serially, with a 100-MB DelayedArray block size for TENxMatrix. In internal testing, we found that an eight-worker TENxMatrix configuration was actually slower than serial execution. dbMatrix, by contrast, used eight DuckDB threads plus eight native threads for the db_svd operator and score projection, with db_svd settings of tol = 1e-5, maxit = 1,000, and ncv = 40. We compared highly variable feature selections across backends using a tie-aware relative cutoff tolerance of 1e^−10^, and verified that principal-component outputs had the expected rank and finite coordinates, loadings, and eigenvalues.

Figure 2e compared dbSequence::pool() against movAPA::subsetPACds(group = “cell_id”, pool = TRUE) on identical prepared bin-level inputs, comparing the same peak-by-cell outputs. Mapping, object creation, and output validation were not timed. For dbSequence::pool(), the full database query was evaluated within the timed call. Output equivalence could only be validated at 1,000 cells, the largest size at which both methods finished. Figure 2f compared GiottoSeq::calculateRUD(method = “smartRUD”, pseudo = 0, min_counts = 0) against movAPA::movAPAindex(method = “smartRUD”, sRUD.oweight = TRUE), using shared proximal and distal annotations and count inputs prepared outside timing. Successful paired outputs through 200,769 cells matched within an absolute tolerance of 1e^−8^. Both benchmarks allowed 600 second per operation and 1,800 second for setup. Figure 2f used a 256GiB R vector and 14GiB RSS limit for the isolated 2.1-million-cell movAPA setup.

## Data Availability

The Visium HD CytAssist colorectal cancer dataset used as the parent for the Figure 2c semisynthetic matrices was obtained from 10x Genomics (https://www.10xgenomics.com/datasets/visium-hd-cytassist-gene-expression-libraries-of-human-crc). Figure 2d used public full-dataset inputs from Stereo-seq mouse embryo cell-bin data (https://ftp.cngb.org/pub/SciRAID/stomics/STDS0000058/stomics/E16.5_E1S3_cell_bin.h5ad), Visium HD 3′ human ovarian cancer segmented outputs (https://cf.10xgenomics.com/samples/spatial-exp/4.0.1/Visium_HD_3prime_Human_Ovarian_Cancer/Visium_HD_3prime_Human_Ovarian_Cancer_segmented_outputs.tar.gz), Visium HD human colon cancer segmented outputs (https://cf.10xgenomics.com/samples/spatial-exp/4.0.1/Visium_HD_Human_Colon_Cancer/Visium_HD_Human_Colon_Cancer_segmented_outputs.tar.gz), and a public CosMX colon processed Seurat object (https://smi-public.objects.liquidweb.services/cosmx_colon_pdr_wtx/sempreqc.RDS).

The Visium HD 3′ ovarian clear cell carcinoma (OCCC) analysis used the public 10x Genomics dataset (https://www.10xgenomics.com/datasets/visium-hd-three-prime-ovarian-cancer-fresh-frozen). Synthetic and semisynthetic benchmark inputs were generated as described in **Methods**. Complete data-generation and validation code is available as described in **Code Availability**.

## Code Availability

All dbverse libraries can be installed at https://github.com/dbverse-org. GiottoDB can be installed and accessed at https://github.com/giotto-suite/GiottoDB. GiottoSeq can be installed and accessed at https://github.com/giotto-suite/GiottoSeq. Benchmarking scripts can be installed and accessed at https://github.com/drieslab/dbverse-paper.

## Acknowledgements

We thank members of the Dries lab for their feedback on this manuscript. Our work was funded by the National Human Genome Research Institute (1R01HG014487-01), the Boston University Rafik B. Hariri Institute for Computing Graduate Student Fellows program, Graduate Education for Minorities, the Boston Medical Center Carter Fund, and the Chan Zuckerberg Initiative’s Essential Open Source Software for Science program.

## Author Contributions

E.C.R. conceived and designed the study; co-developed the dbverse architecture; implemented dbverse, GiottoDB and GiottoSeq; performed the computational analyses and software validation; prepared the figures; and wrote and revised the manuscript. V.J. contributed to GiottoDB development, performed the Visium HD 3′ ovarian cancer analysis and assisted with figure preparation. J.C. contributed to GiottoDB development. T.R. contributed to GiottoDB development and assisted with figure preparation. I.A. contributed to GiottoDB development. R.D. supervised study conception, dbverse architecture and methodology; interpreted the results; and revised the manuscript.

## Supplementary Figures

**Supplementary Fig. 1.**
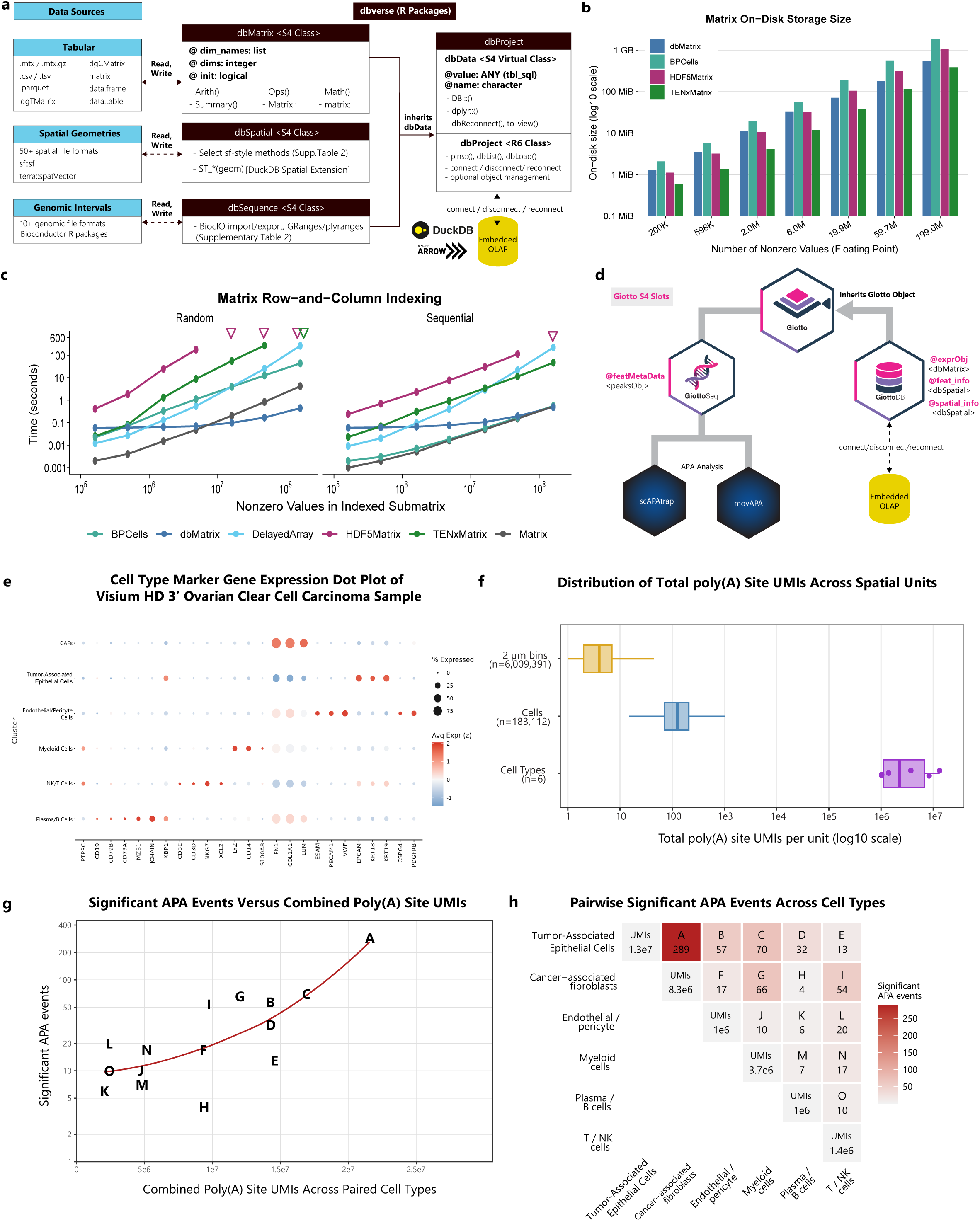
dbverse architecture, matrix benchmarks and APA analyses in Visium HD 3′ data. **a,** Diagram showing dbverse architecture linking tabular, spatial, and genomic data sources to dbMatrix, dbSpatial, dbSequence and dbProject within an embedded OLAP database, respectively. **b,** Barplot showing on-disk storage footprint across matrix backends for increasing matrix sizes at fixed 99% sparsity. **c,** Line plots showing runtimes for random and sequential row-and-column indexing followed by colSums to force computation and value return. ‘▽’ denotes a 600-s timeout or an operation not attempted after a preceding setup timeout; ‘×’ denotes an out-of-memory failure. Successful points show the median of five runs per method and dataset size. **d,** Diagram showing GiottoDB integration with dbverse-backed representations, GiottoSeq APA functionality, scAPAtrap and movAPA interoperability, and the embedded OLAP engine. **e,** Dot plot showing cell-type marker expression. Point size indicates the percentage of cells expressing each gene and color indicates scaled mean expression. **f,** Distribution of total poly(A)-site UMI counts per spatial unit across 2µm bins, cells and cell-type aggregates. **g,** Scatterplot showing the relationship between combined poly(A) site UMI read depth and significant APA events across pairwise major-cell-type contrasts. The red curve indicates a LOESS trend; letters A-O identify the contrasts shown in h. **h,** Heatmap of pairwise major-cell-type contrasts. Diagonal entries show total poly(A) site UMI read depth and upper-triangle entries show significant-event counts.

## Supplementary Tables

**Supplementary Table 1 | dbverse input and interoperability support. Supported file formats, in-memory R object classes and conversion paths for dbMatrix, dbSpatial, dbSequence, and dbProject.**

Supplementary Tables 1 and 2 spreadsheet

**Supplementary Table 2 | Database-backed operations implemented in dbverse. Matrix, spatial and genomic operations supported by dbverse libraries, including corresponding R package APIs where applicable.**

Supplementary Tables 1 and 2 spreadsheet

**Supplementary Table 3 | Additional dbverse operation benchmarks. Runtime benchmarks for additional matrix, spatial and genomic operations comparing dbverse to commonly used in-memory and file-backed R packages.**

Supplementary Table 3 data file

Supplementary Table 3 figure

**Extended Data Table 1 | Source data for Fig. 1a. Published omics dataset metadata and estimated count matrix footprints used to plot memory footprint over time.**

Extended Data Table 1 data file

